# A probabilistic atlas of the human thalamic reticular nucleus derived from 7T MRI

**DOI:** 10.64898/2026.06.22.733673

**Authors:** Zuzanna Kotwicka, Omer Faruk Gulban, Logan Dowdle, Ryszard Auksztulewicz, Michelle Moerel

## Abstract

The thalamic reticular nucleus (TRN) is a thin, inhibitory shell surrounding the thalamus. It regulates the thalamocortical information flow, and thereby plays a central role in attention, task switching, and the sleep-wake cycle. Despite its importance, the TRN remains poorly studied in the human brain. This is largely because its small size and deep anatomical location limit its visibility with conventional non-invasive neuroimaging techniques. Here, we assessed whether the human TRN can be reliably visualised and segmented *in vivo* using ultra-high field (UHF) magnetic resonance imaging (MRI) at 7 Tesla. High resolution (0.35 mm isotropic) partial-brain T_2_* and T_1_ scans were acquired from healthy individuals, followed by manual delineation of the TRN. These *in vivo* segmentations were compared with TRN estimates obtained from two high-quality postmortem datasets serving as an anatomical reference. *In vivo* segmentations of TRN volume and thickness closely matched measurements derived from the postmortem reference datasets, and quantitative comparisons showed high consistency in TRN shape and location across individuals while also capturing meaningful inter-individual variability. Using these segmentations, we constructed a publicly available probabilistic atlas of the human TRN. This atlas provides a new resource for incorporating TRN anatomy into functional, structural, and clinical neuroimaging studies. Our findings demonstrate that the human TRN can be robustly mapped *in vivo* at 7T and establish a foundation for future investigations into its structure and function.

## 1. Introduction

The thalamus is a central hub of the human brain, routing and shaping sensory, motor, and behavioural information before it reaches the cortex (Shine et al., 2023). Surrounding the thalamus lies the thalamic reticular nucleus (TRN), a thin sheet of GABAergic neurons that plays an essential role in regulating thalamocortical communication (Li et al., 2020). The TRN is organised in modality-specific sections that interface with distinct thalamic and cortical regions (Guillery et al., 1998; Li et al., 2020), enabling it to influence processing across multiple sensory and cognitive domains. Positioned strategically between the thalamus and the cortex, the TRN modulates the flow of information by selectively enhancing or suppressing thalamic output, thereby influencing attention, sensory gating, and transitions between sleep and wakefulness (Takata, 2019). Because of these roles, TRN dysfunction has been implicated in a range of clinical conditions. Altered TRN-mediated inhibition is hypothesized to contribute to tinnitus (Henton & Tzounopoulos, 2021; Rauschecker et al., 2010), sleep disturbances (Sun et al., 2025; Vantomme et al., 2019), and the positive, negative, and cognitive symptoms of schizophrenia (El Khoueiry et al., 2022; Pratt & Morris, 2015; Zhu et al., 2021). Disruptions in TRN connectivity have also been proposed as a mechanism underlying attentional deficits in various psychiatric disorders (Madureira et al., 2010; Zikopoulos & Barbas, 2012).

Despite its widespread functional relevance, the TRN remains poorly characterised in the human brain. Most existing knowledge comes from rodent models, which differ substantially from humans in TRN size, internal organisation (Xu et al., 2022), and thalamocortical connectivity (Brown & Bowman, 2002; Schaeffer et al., 2020), limiting direct translational interpretation. This gap in our understanding largely stems from methodological limitations: the TRN’s small size and deep location make it difficult to visualise using conventional non-invasive imaging methods (Keun et al., 2021). Ultra-high field (UHF) magnetic resonance imaging (MRI) at 7 Tesla offers a potential solution. Compared with clinical field strengths, UHF MRI offers increased signal-to-noise ratio that can be leveraged to increase spatial resolution (Yacoub et al., 2001). This has proven highly valuable for imaging small subcortical nuclei (Bazin et al., 2020; Denison et al., 2014; Isaacs et al., 2020; Moerel et al., 2015; Müller-Axt et al., 2021; Patriat et al., 2023; Sitek et al., 2019).

Here, we use high-resolution 7T MRI to determine whether the TRN can be consistently visualised and manually segmented *in vivo*. We compare these segmentations to high-quality postmortem delineations (Alkemade et al., 2022) to evaluate anatomical correspondence, and assess variability in TRN shape, location, and morphology across individuals. Building on these results, we construct a probabilistic atlas of the human TRN in standard MNI space and make it publicly available. This resource is intended to support future studies of TRN structure and function, including for those without access to ultra-high field imaging or the time resources required for manual segmentation. As such, the atlas offers a tool for integrating TRN anatomy into functional, structural, and clinical investigations, representing a first step toward the systematic characterisation of the TRN in human neuroscience.

## 2. Methods

### 2.1. Participants

*In vivo* data were obtained from two sources. First, we used a publicly available 7T MRI dataset from Gulban et al. (2022), which provides high-resolution, quantitative structural scans from healthy young adults (2 females and 3 males, n = 5), aged from 27 to 35. From this dataset, we selected four participants.

Second, we acquired new high-resolution 7T MRI data from eleven healthy individuals (8 females and 3 males, n = 11), aged from 19 to 25. Participants were screened for standard contraindications to 7T MRI and excluded if they reported a history of neurological or psychiatric disorders, or uncorrectable visual impairments. Of these newly collected datasets, the six datasets with the highest image quality for TRN segmentation were included in the analysis. Datasets of the remaining five subjects were excluded due to their incomplete nature (n = 2) and motion-related artefacts (n = 3) which hindered TRN visibility. All procedures for data collection were approved by the Ethical Review Committee of the Faculty of Psychology and Neuroscience (ERCPN), Maastricht University, the Netherlands (ERCPN-263_12_02_2023). All participants provided written informed consent before participation, and a short debriefing was provided at the end of the study.

In addition, we used a publicly available postmortem dataset (Alkemade et al., 2022), consisting of two human brains (pm-1 and pm-2; ages at death: 75 and 59 years) for which both MRI and detailed histological imaging data are available. Using the histological imaging data (specifically the ‘block face’ images), we performed precise manual TRN segmentations in each brain. These postmortem segmentations served as an anatomical reference for validating the *in vivo* TRN delineations.

### 2.2. MRI data acquisition

MRI data were collected using a Siemens MAGNETOM 7 Tesla MRI scanner with a 1Tx/32Rx-channel head coil at Scannexus (Maastricht, the Netherlands). Each scanning session lasted approximately 60 minutes, during which multiple *in vivo* anatomical datasets were collected.

Data acquisition followed the protocol described by Gulban et al. (2022) and included T_1_ and T_2_* imaging at high spatial resolution (see Table 1 for acquisition parameters). Specifically, a Magnetization Prepared 2 Rapid Gradient Echoes (MP2RAGE) sequence was used for collecting quantitative T_1_ maps. Whole-brain anatomical images were acquired at 0.7 mm isotropic resolution (repetition time [TR] = 5000 ms, echo time [TE] = 2.47 ms, inversion time [TI_1,2_] = [900, 2750] ms, flip angles [FA_1,2_] = [5°, 3°], field of view [FOV] = 224 mm, GRAPPA = 3, partial Fourier = 6/8, bandwidth = 250 Hz/pixel). In addition, a slightly modified MP2RAGE sequence was used to acquire partial brain scans covering a horizontal slab at a higher resolution of 0.35 mm isotropic (TR = 5000 ms, TE = 3.45 ms, TI_1,2_ = [800, 2700] ms, FA_1,2_ = [4°, 5°], slab dimensions = 576 x 576 x 120 voxels, FOV = 201.6 x 201.6 x 42 mm, GRAPPA = 2, partial Fourier = 6/8, bandwidth = 250 Hz/pixel) (Table 1). The partial brain scan was acquired twice. The slab was positioned to include the thalamus, basal ganglia and surrounding subcortical structures.

**Table 1.** Acquisition parameters for scanning protocols.

| Sequence | Partial brain ME GRE | Partial brain MP2RAGE | Whole brain MP2RAGE |
| --- | --- | --- | --- |
| Voxel size | 0.35 x 0.35 x 0.35 mm<sup>3</sup> | 0.35 x 0.35 x 0.35 mm<sup>3</sup> | 0.7 x 0.7 x 0.7 mm<sup>3</sup> |
| Field of view (FOV) | 201.6 x 201.6 x 36.4 mm | 201.6 x 201.6 x 42 mm | 224 mm |
| Slab dimensions | 576 x 576 x 120*<br>576 x 576 x 104** | 576 x 576 x 120 | N/A |
| Bandwidth | 310 Hz/Pixel*<br>240 Hz/Pixel** | 250 Hz/Pixel | 250 Hz/Pixel |
| Repetition time (TR) | 32 ms*<br>30 ms** | 5000 ms | 5000 ms |
| Echo time (TE) | [3.60, 7.20, 10.80, 14.40, 18, 21.60] ms*<br>[3.83, 8.20, 12.57, 16.94, 21.31, 25.68] ms** | 3.45 ms*<br>2.91 ms** | 2.47 ms |
| Flip angles (FA) | 12°*<br>11°** | [4°, 5°] | [5°, 3°] |
| Inversion time (TI) | N/A | [800, 2700] ms | [900, 2750] ms |
| GRAPPA | 2 | 2 | 3 |
| Partial Fourier | Off | 6/8 | 6/8 |
| Acquisition time | 15 min*<br>11 min** | 10 min | 8 min |

\*our dataset
\*\*dataset from (Gulban et al., 2022)

Partial brain T_2_* data, covering the same brain structures as the partial brain MP2RAGE dataset, were acquired using a multi-echo gradient-recalled echo (ME GRE) sequence. Four repetitions were collected at 0.35 mm isotropic resolution, acquired twice per phase-encode direction (anterior-posterior and right-left). Acquisition parameters included TR = 32 ms, TE_1-6_ = [3.60, 7.20, 10.80, 14.40, 18, 21.60] ms, FA = 12°, slab dimensions = 576 x 576 x 120 voxels, FOV = 201.6 × 201.6 × 36.4 mm, GRAPPA = 2, bandwidth = 310 Hz/pixel (see Table 1).

### 2.3. Data processing

The raw data obtained from the scanner were first converted from DICOM to NIfTI format using MRIcroGL to enable further processing. The two dataset types (MP2RAGE and ME GRE) were processed through separate preprocessing pipelines (see Fig. 1 for a pipeline overview), implemented with a combination of Python scripts from Gulban et al. (2022) and ITK-SNAP. For the ME GRE data, preprocessing began with noise reduction using the Transform domain NOise Reduction with DIstribution Corrected principal component analysis (tNORDIC) method (Ponticorvo et al., 2024). tNORDIC, which improves signal quality by attenuating hardware-related noise, was implemented using custom MatLab scripts. Subsequently, the images were cropped to reduce file size and to retain only the region of interest (ROI), defined as the volume spanning from the anterior to posterior horns of the lateral ventricles, including the thalamus and surrounding structures, such as the internal capsule and the basal ganglia. Next, the 6 echoes were averaged into one image per repetition and those images were upsampled to an isotropic resolution of 0.175 mm (using cubic interpolation) to ensure consistent voxel geometry and improve the accuracy of subsequent registration and ROI-based analyses. Then, motion correction was applied across all repetitions using Greedy (Yushkevich et al., 2016; available at https://github.com/pyushkevich/greedy); with rigid body registration (6 degrees of freedom), normalised cross correlation (NCC, radius 2×2×2), and multi-resolution optimisation scheme (100×50×10 iterations), during which repetitions 2, 3, and 4 were registered to repetition 1. To prioritise alignment within the ROI, a 3D mask covering the thalamus was used during coregistration. Next, echoes in the original cropped images were split, upsampled to an isotropic resolution of 0.175 mm, and motion corrected by applying the previously computed coregistration matrix to repetition 1. Applying the same transformations avoids redundant interpolation and ensures consistency with the earlier motion correction step. The upsampled, motion corrected single echo images were then merged back together. All ME GRE images acquired across the same axis were combined into one averaged file and a composite image was generated. At this stage, a SimpleITK bias field correction was implemented into the pipeline to remove bias field-induced intensity variations visible on the images (Kanakaraj et al., 2024). Although such correction is not standard practice for quantitative T_2_* mapping because it introduces data-driven intensity scaling that can affect absolute parameter values, this step was included as it substantially improved the visibility of the TRN. After bias field correction, where non-decaying artefacts were identified and corrected, we fitted a voxel-wise mono-exponential decay model to the ME GRE data across the six echoes. This was not done to obtain quantitative T_2_* values, but to combine the multi-echo data into a single image. Here, the exponential fit is preferential to simple averaging across echoes, as it suppresses the contribution from later, lower-SNR echoes. The resulting images are referred to as T_2_w maps, reflecting the T_2_-weighted contrast rather than quantitative T_2_* estimates.

**Figure 1.**
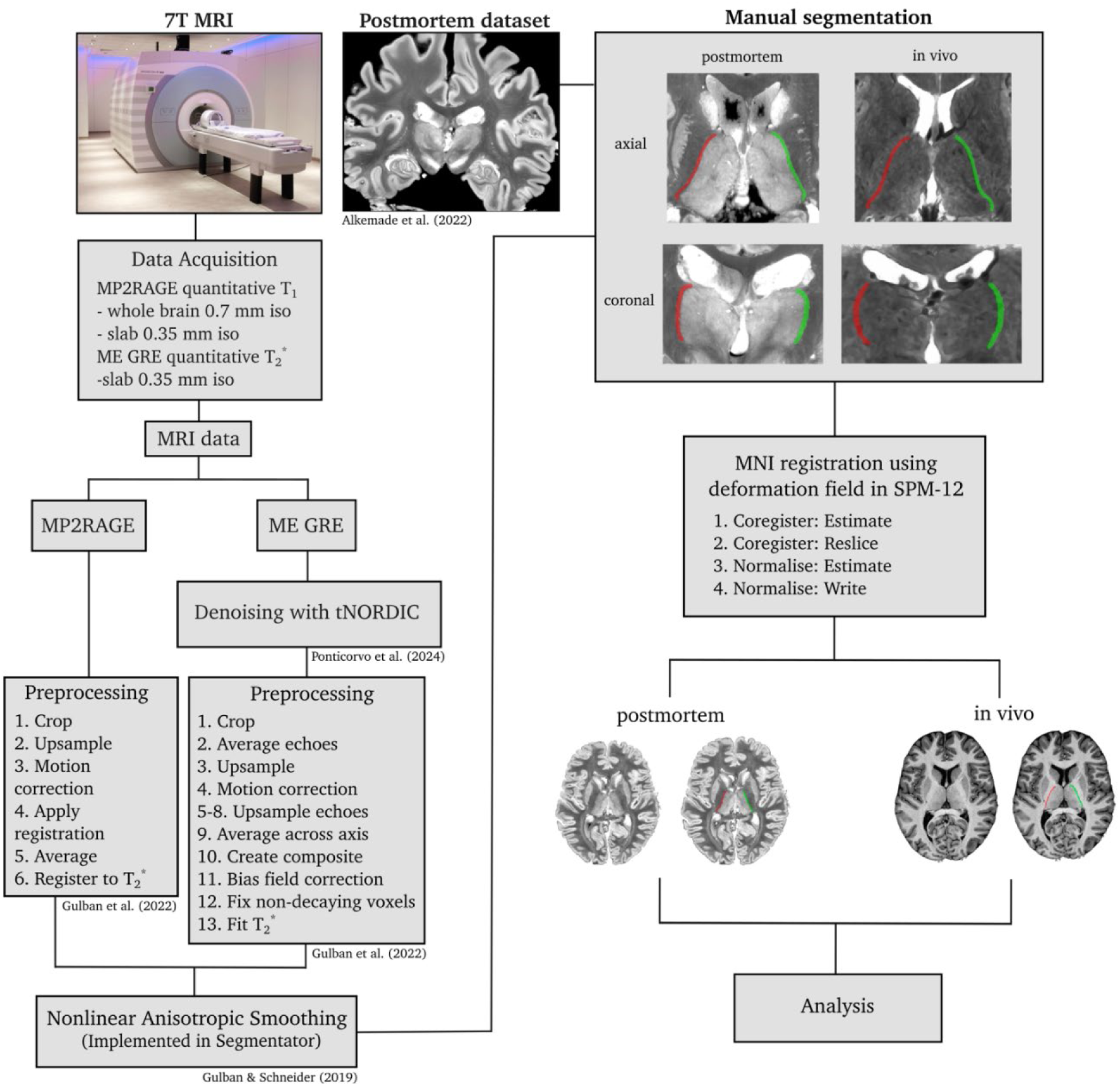
Pipeline Overview. Flowchart illustrating the pipeline for visualisation and segmentation of the TRN in *in vivo* and postmortem datasets. *In vivo* images were acquired using ultra-high field 7T MRI with sequences optimised for subcortical contrast and subsequently denoised. Preprocessing steps, including upsampling, motion correction, and bias field correction were applied to enhance TRN contrast. Manual segmentation was performed for both *in vivo* and postmortem images. Segmentations were aligned to MNI space to reduce inter-individual differences in brain orientation and size, enabling between-subject comparisons.

The MP2RAGE partial-brain data were first cropped to the same ROI as defined for the ME GRE data, and then upsampled to an isotropic resolution of 0.175 mm (using cubic interpolation). Head motion was corrected by registering repetition 2 to the first scan (repetition 1) using a 3D mask. The motion-corrected repetitions were then averaged and registered to the T * image space using an affine transformation manually defined in ITK-SNAP and applied with Greedy (Yushkevich et al., 2016; available at https://github.com/pyushkevich/greedy) using linear interpolation. Finally, nonlinear anisotropic smoothing was performed on both preprocessed partial-brain datasets using the Segmentator tool (Gulban & Schneider, 2019), applying the CURED smoothing type with noise and feature scales set to 1, and 10 iterations, saving every 5^th^ iteration. The effects of denoising, bias correction, and smoothing are shown in Supplementary Fig. 1.

The MP2RAGE whole-brain images were defaced using the Segmentation and ImCalc tools in the Statistical Parametric Mapping toolbox (SPM12) for Matlab. No other preprocessing steps were performed on either the MP2RAGE or postmortem datasets.

### 2.4. Manual delineations

Following preprocessing, the TRN was manually segmented in ITK-SNAP, resulting in binary masks for each participant and hemisphere. Segmentation was performed manually in native space using both the T_1_ and T_2_*w images for the *in vivo* datasets, and the block face postmortem images. For the *in vivo* data, delineations were primarily guided by the T_2_*w images, with UNI T_1_ images consulted as complementary anatomical references. All segmentations were carried out using a WACOM Cintiq 16 drawing tablet and were guided by local image contrast (with the TRN appearing lighter than adjacent structures in T_2_*w images) and anatomical landmarks, including the thalamus and internal capsule.

When completed, binary morphological smoothing was applied to each segmentation. To enable between-subject comparisons, all segmentations were brought to the MNI IC6BM152 2009b space (Fonov et al., 2009). This was implemented by registering each participant’s whole brain MP2RAGE image to the MNI template, generating non-linear deformation fields in SPM12. These deformation fields were then applied to the corresponding TRN segmentations, thereby bringing them into MNI space. The normalised segmentations were downsampled to 0.7 mm isotropic using nearest-neighbour interpolation (see Fig. 1 for an overview of the processing pipeline).

### 2.5. Morphological quantification and spatial overlap analysis

The morphology of each TRN segmentation was quantified in native space by measuring its volume and thickness. Volume was calculated as the total number of voxels within the segmented region multiplied by the voxel volume and obtained using the Statistics and Volumes tool in ITK-SNAP. TRN thickness was estimated by performing multiple radial distance measurements between opposing TRN boundaries in both coronal and axial slices using the ITK-SNAP ruler tool. These measurements were averaged to provide a single thickness estimate for each hemisphere.

To assess the anatomical variability and the reliability of manual segmentations, the spatial overlap between the segmentations was quantified using the Dice similarity coefficient, which measures the relative volumetric overlap between two segmentations (eq. 1):

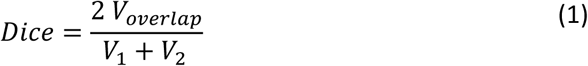

where *V_overlap_* denotes the shared volume between two segmentations, and *V*_1_ and *V*_2_ their respective volumes. To calculate the relative overlap of segmentations between hemispheres, a custom MATLAB script was used to mirror the brain across the sagittal midline. Then, the Dice coefficient was calculated based on the proportion of overlapping volume between the left and right hemisphere TRN segmentations, with higher values signifying higher overlap (1 = full overlap, 0 = no overlap). For between-subject analyses, Dice scores were computed for all possible pairs of segmentations, including those derived from the postmortem datasets. In addition to overlap-based metrics, shape similarity was quantified using the Average Hausdorff Distance (AHD), which captures the average surface-to-surface distance between two segmentations (eqs. 2, 3):

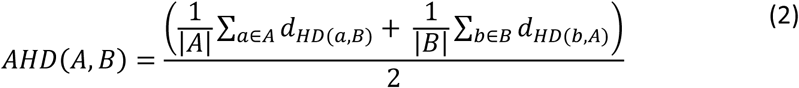

where A and B represent the sets of boundary voxels of two segmentations, and

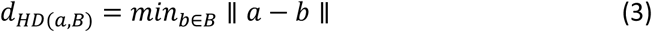

The AHD reflects the mean of the minimum distances between segmentation boundaries in both directions. This metric thus provides a complementary measure to overlap-based measures, as it is sensitive to discrepancies in both spatial location and shape (i.e., boundary discrepancies; Taha & Hanbury, 2015). AHD calculations were performed in MATLAB, both within and between datasets, with all possible *in vivo* and postmortem-based segmentation pairs included in the between-dataset analysis. All Dice and AHD calculations were performed in MNI space.

Finally, a probabilistic map of the human TRN was created by summing all individual *in vivo* segmentations using a custom MATLAB code. Voxel intensities in the probabilistic map ranged from 0 to 10, where 0 indicates the absence of a TRN across all individual datasets, and 10 indicates locations where the TRN was present in all ten individual *in vivo* segmentations.

## 3. Results

### 3.1. Data quality and preprocessing

Data quality was evaluated after each preprocessing step to assess its impact on the visibility of the TRN and surrounding subcortical structures (Supplementary Fig. 1). Initial processing of the ME GRE data with tNORDIC (Ponticorvo et al., 2024) resulted in substantial noise reduction, observed as a decrease in high-frequency fluctuations in the images. Subsequent bias field correction further increased the TRN contrast, improving its visibility and discriminability from surrounding structures. Alignment of repeated dataset acquisitions required manual corrections, as the automatic alignment procedure frequently failed. This failure was likely caused by the substantial cropping of the data, which reduced the number and extent of high contrast areas that typically serve as reference points for registration. Finally, spatial smoothing enhanced the signal to noise ratio and further increased the tissue contrast (Supplementary Fig. 1).

### 3.2. Manual segmentation

Following preprocessing, the TRN was identifiable on the T_2_*w images in 10 out of 15 *in vivo* datasets. For simplicity, these 10 subjects are referred to below as subjects 1-10 (sub-1 to sub-10).

On the *in vivo* T_2_*w images, the TRN consistently appeared as a light grey, thin, elongated structure encapsulating the darker grey thalamus and surrounded by the internal capsule (Fig. 2A). In the postmortem block face data, the TRN was visible even more clearly, with distinct dark grey boundaries separating it from the thalamus (Fig. 2A). Across both *in vivo* and postmortem datasets, the TRN appearance varied with the plane and level of section. Specifically, the TRN appeared more rounded and compact in superior and rostral regions, and increasingly elongated and curled in inferior and caudal regions (Fig. 2B).

**Figure 2.**
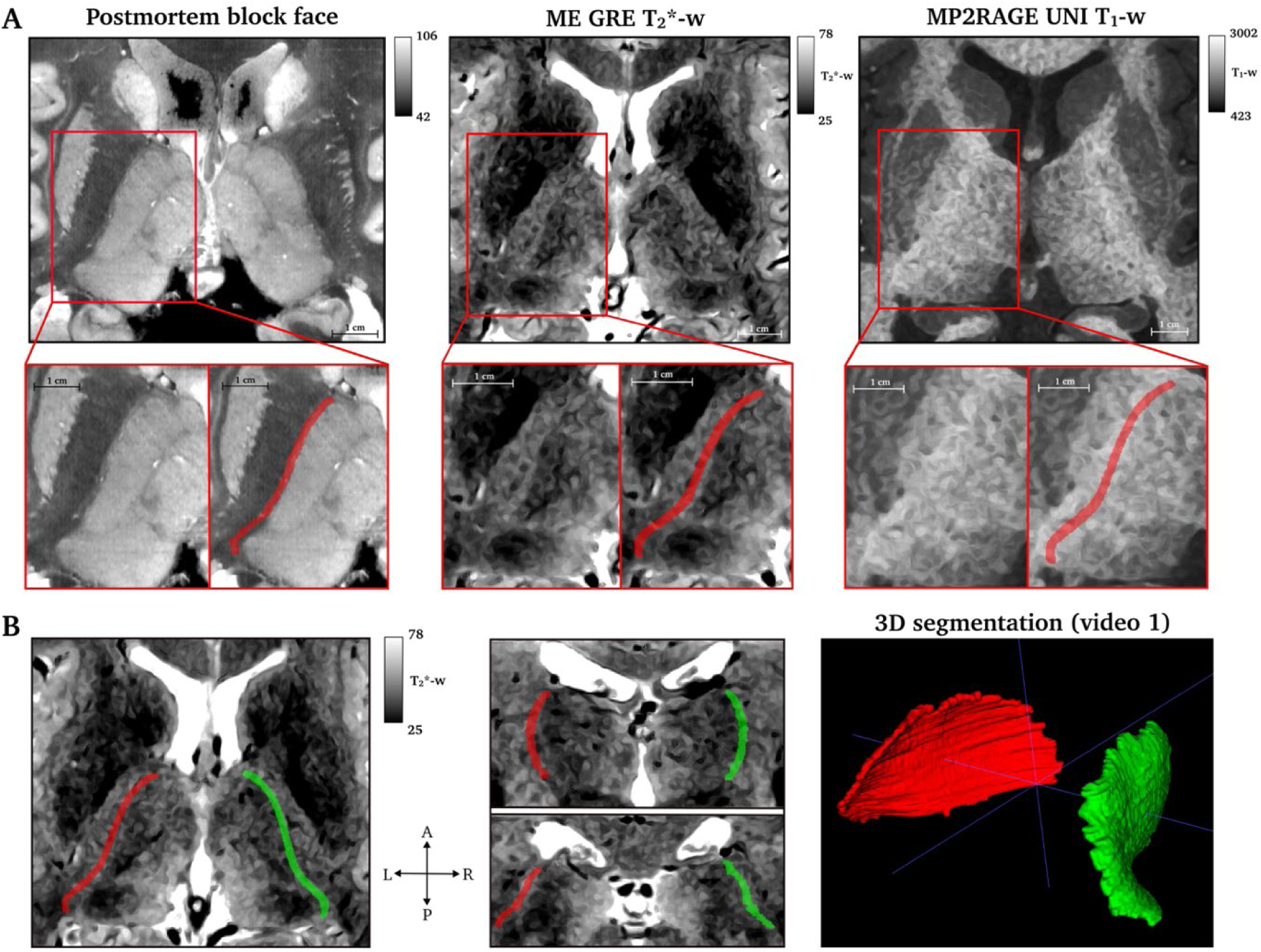
Data quality. (A) Visibility of the TRN in a block face postmortem dataset (left), and in a representative subject (sub-9) shown on smoothed *in vivo* ME GRE weighted T * (T *-w; middle) and smoothed *in vivo* MP2RAGE UNI weighted T_1_ (T_1_-w; right) images. In the zoomed view, the left TRN appears as an elongated arch in a lighter colour surrounding the thalamus. The corresponding TRN segmentation is overlayed in red. (B) Visualisation of the TRN segmentation on axial (left) and coronal (middle) slices, with the upper panel being more anterior, displayed as red and green labels. The right panel shows a representative frame from a 3D rendering of the TRN segmentation (Supplementary Video 1; available on https://doi.org/10.5281/zenodo.20446658). A = anterior, P = posterior, R = right, L = left.

### 3.3. TRN morphology

Morphological quantification of the manual TRN segmentations showed relatively stable volumes and thicknesses both within and across datasets (Table 2, Fig. 3D-E). TRN volumes ranged from 562 mm^3^ (in pm-1, right TRN) to 775 mm^3^ (in sub-2, right TRN) across all datasets, with an average volume of 683.5 mm^3^ (standard deviation [SD] = 65.4 mm^3^) and no volumes exceeding ± 2 SD from the mean. No hemispheric differences were observed (mean [SD] right TRN = 687.2 [64.3] mm^3^, mean [SD] left TRN = 679.8 [69.2] mm^3^; paired t-test: t(11) = 0.835, p = 0.422; Fig. 3D). Volume ranges of the *in vivo* and postmortem-based segmentations were similar, with histological data tending toward the lower end of the distribution (Fig. 3D). Consequently, the average volume derived from *in vivo* data alone was slightly higher (mean = 698.8 mm^3^, ranging from 567 mm^3^ in the left TRN of sub-10 to 775 mm^3^ in the right TRN of sub-2) than that derived from the postmortem data (mean = 607.3 mm^3^, ranging from 562 mm^3^ in the right TRN of pm-1 to 652 mm^3^ in the left TRN of pm-2). A similarly consistent pattern was observed for thickness measurements, with no statistically significant differences between hemispheres (mean [SD] right TRN = 1.333 [0.108] mm, mean [SD] left TRN = 1.282 [0.076] mm; paired t-test: t(11) = 1.790, p = 0.101; Fig. 3E), or between the *in vivo* and postmortem data. The average thickness across all segmentations was 1.307 mm (SD = 0.095), ranging from 1.164 mm in pm-2 (right TRN) to 1.489 mm in sub-9 (right TRN). No significant correlation between volumes and thicknesses was observed (right TRN: r(10) = 0.241, p = 0.450; left TRN: r(10) = −0.067, p = 0.837; Fig. 3A-C).

**Figure 3.**
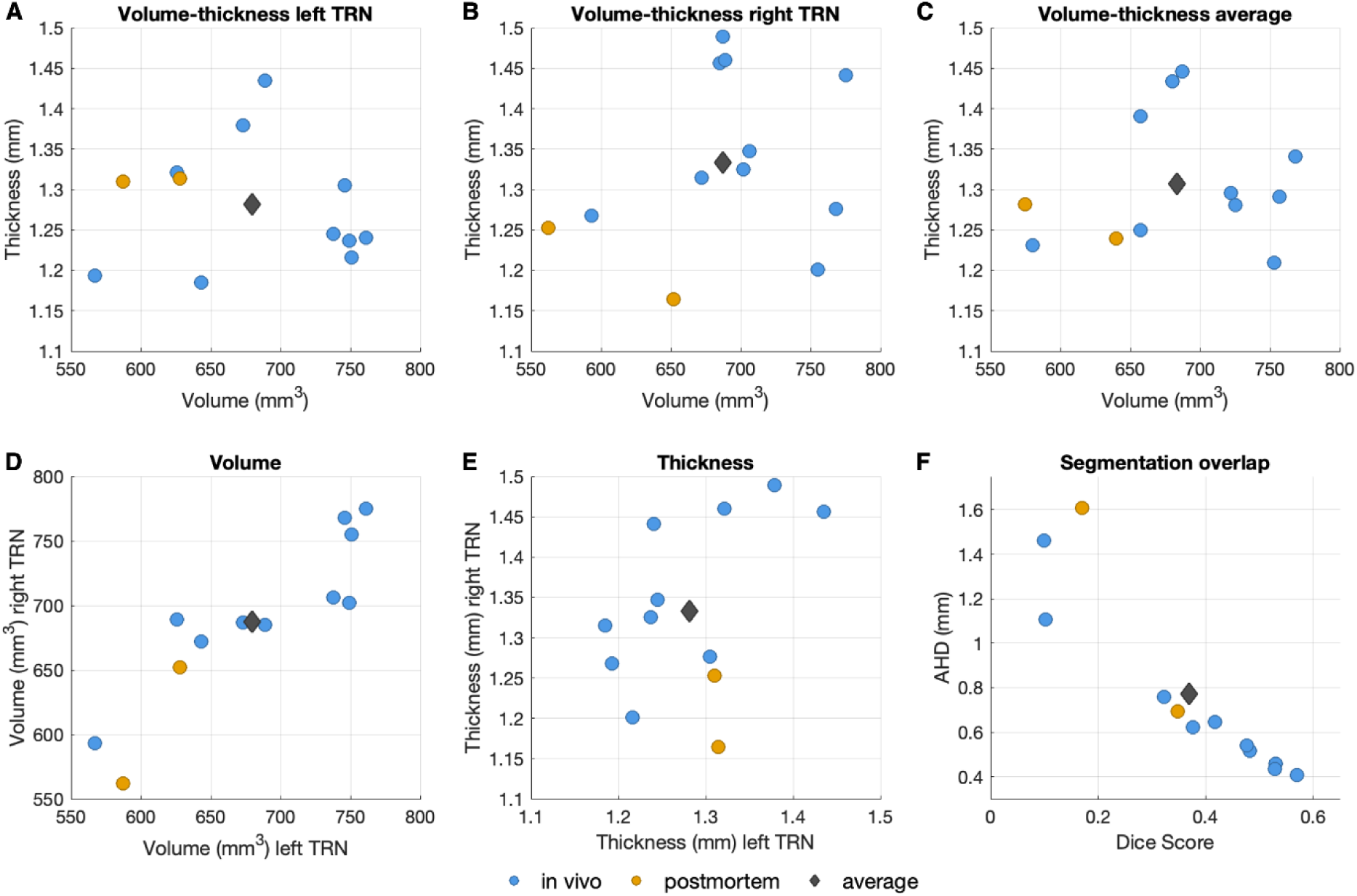
TRN morphometry and segmentation overlap across datasets. (A-C) Relationship between TRN volume (in mm^3^) and thickness (in mm) for the left hemisphere (A), right hemisphere (B), and averaged across hemispheres (C). (D) Comparison of left and right TRN volume. (E) Comparison of left and right TRN thickness. (F) Within-dataset hemisphere overlap quantified as Dice coefficients (x-axis) and AHD values (y-axis), where higher Dice and lower AHD values represent higher overlap. Across panels, blue circles represent individual *in vivo* datasets, orange circles represent postmortem datasets, and the grey diamond indicates the average across segmentations.

**Table 2.** TRN volume and thickness.

| Subject | Hemisphere | Volume (mm<sup>3</sup>) | Thickness (mm) | Volume average (mm<sup>3</sup>) | Thickness average (mm) |
| --- | --- | --- | --- | --- | --- |
| Sub-1 | Right | 706 | 1.347 | 722.0 | 1.296 |
|  | Left | 738 | 1.245 |  |  |
| Sub-2 | Right | 775 | 1.441 | 768.0 | 1.341 |
|  | Left | 761 | 1.240 |  |  |
| Sub-3 | Right | 768 | 1.276 | 757.0 | 1.291 |
|  | Left | 746 | 1.305 |  |  |
| Sub-4 | Right | 685 | 1.456 | 687.0 | 1.446 |
|  | Left | 689 | 1.435 |  |  |
| Sub-5 | Right | 702 | 1.325 | 725.5 | 1.281 |
|  | Left | 749 | 1.237 |  |  |
| Sub-6 | Right | 689 | 1.460 | 657.5 | 1.391 |
|  | Left | 626 | 1.321 |  |  |
| Sub-7 | Right | 755 | 1.201 | 753.0 | 1.209 |
|  | Left | 751 | 1.216 |  |  |
| Sub-8 | Right | 672 | 1.315 | 657.5 | 1.250 |
|  | Left | 643 | 1.185 |  |  |
| Sub-9 | Right | 687 | 1.489 | 680.0 | 1.434 |
|  | Left | 673 | 1.379 |  |  |
| Sub-10 | Right | 593 | 1.268 | 580.0 | 1.231 |
|  | Left | 567 | 1.193 |  |  |
| Pm-1 | Right | 562 | 1.253 | 574.5 | 1.282 |
|  | Left | 587 | 1.310 |  |  |
| Pm-2 | Right | 652 | 1.164 | 640.0 | 1.239 |
|  | Left | 628 | 1.314 |  |  |
|  |  |  |  | 683.5 | 1.307 |

### 3.4. Within-dataset quantification of shape similarity and overlap

Morphological quantification of within-dataset TRN shape similarity and overlap revealed subtle but meaningful hemispheric differences in TRN anatomy (Table 3; Fig. 3F). Overall, results showed moderate volumetric overlap between hemispheres, with Dice coefficients ranging from 0.100 (sub-8) to 0.570 (sub-5). The mean Dice coefficient was 0.369 (SD = 0.166) across all segmentations and 0.391 (SD = 0.170) when considering the *in vivo* segmentations only.

**Table 3.** Within-dataset hemispheric Dice coefficients and Average Hausdorff Distances (AHD)

| Dataset | Dice | AHD (mm) |
| --- | --- | --- |
| Sub-1 | 0.530 | 0.457 |
| Sub-2 | 0.323 | 0.758 |
| Sub-3 | 0.483 | 0.516 |
| Sub-4 | 0.103 | 1.108 |
| Sub-5 | 0.529 | 0.434 |
| Sub-6 | 0.570 | 0.406 |
| Sub-7 | 0.417 | 0.645 |
| Sub-8 | 0.100 | 1.462 |
| Sub-9 | 0.477 | 0.539 |
| Sub-10 | 0.377 | 0.622 |
| Pm-1 | 0.348 | 0.693 |
| Pm-2 | 0.170 | 1.607 |
| Average | 0.369 | 0.771 |

Despite these moderate overlap values, AHD values were relatively low, indicating high hemispheric symmetry in both shape and anatomical location. Mean AHD values were 0.771 mm (SD = 0.404 mm) for all segmentations and 0.695 mm (SD = 0.339 mm) for the *in vivo* subjects. Within dataset TRN overlap as assessed by Dice coefficient vs. AHD showed a significant negative correlation (Fig. 3F; *r*(10) = −0.918, *p* = 2.58 x 10^-5^), confirming that they captured convergent information.

Within the *in vivo* cohort, sub-4 and sub-8 represented notable outliers, with elevated AHD values of 1.108 mm and 1.462 mm, respectively, accompanied by low Dice scores (0.103 for sub-4, and 0.100 for sub-8). Similarly, one postmortem dataset (pm-2) showed an AHD value of 1.607 mm together with a low Dice coefficient (0.170). Visual inspection of the corresponding segmentations confirmed that these values reflected genuine hemispheric asymmetry rather than segmentation error, with systematic differences in ventricle size and thalamic position and shape between hemispheres (see Supplementary Fig. 2 for all individual segmentations).

### 3.5. Between-subject quantification of shape similarity and overlap

The between-subject analysis showed lower TRN similarity compared to within-dataset comparisons (Table 4). Across all datasets, the average Dice coefficient was 0.235 (SD = 0.115) and the average AHD was 1.551 mm (SD = 0.679 mm). The highest alignment was observed between sub-5 and sub-7 with a Dice score of 0.423 and AHD score of 0.629 mm. Similarly high correspondence was found for sub-5 and sub-10 (Dice = 0.417, AHD = 0.900 mm). In contrast, the lowest anatomical correspondence within the *in vivo* subjects was observed for sub-8 and sub-10, with a Dice coefficient of 0.175 and an AHD of 1.605 mm.

**Table 4.**
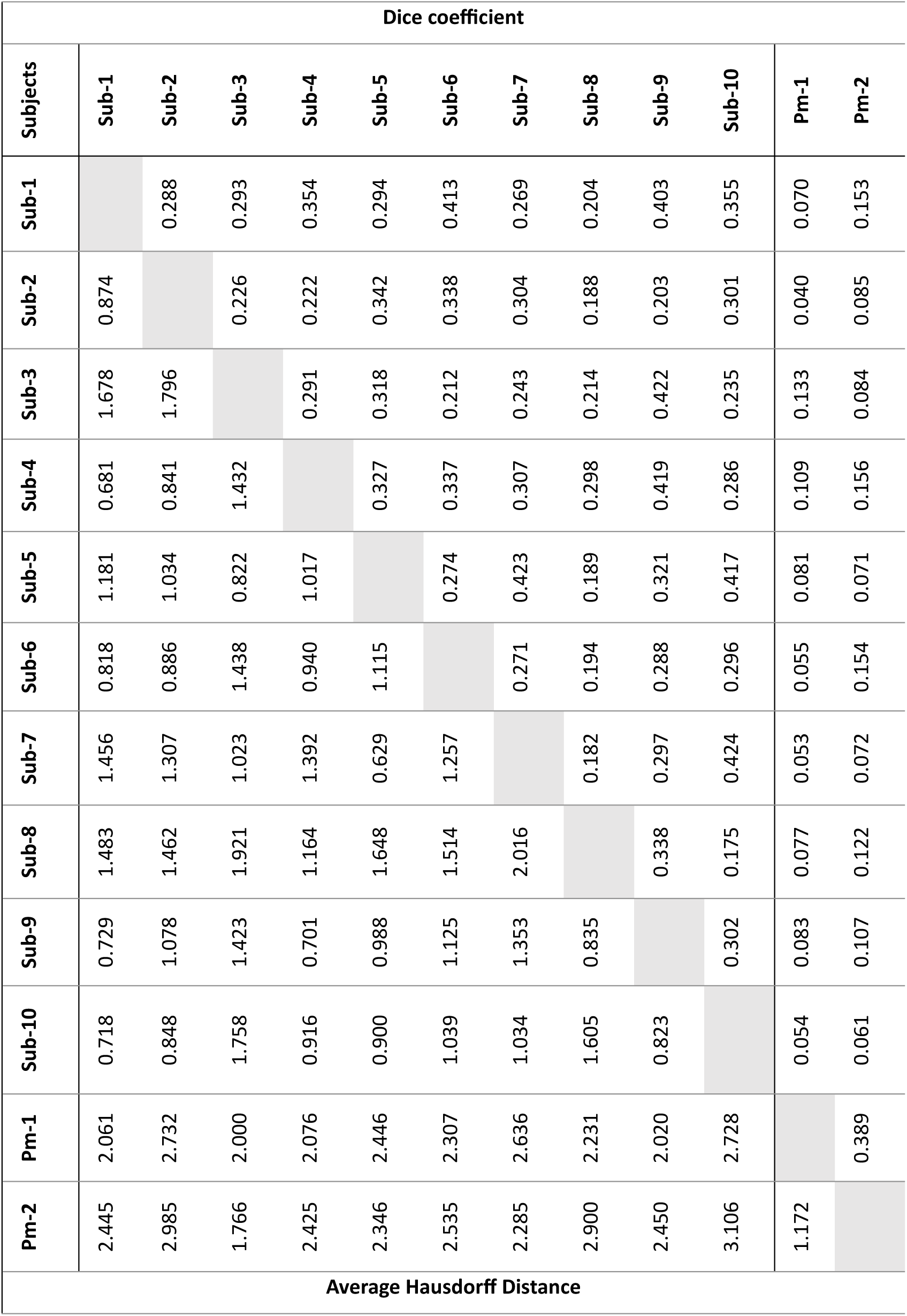
Between-dataset hemispheric Dice coefficients and Average Hausdorff Distances (AHD)

Fig. 4 shows a visual overlay of representative *in vivo* TRN segmentations, illustrating the range of Dice and AHD values observed across pairs. The figure shows both similarities in TRN shape and location, as well as substantial variability across individuals. For example, sub-5 and sub-7, which showed the highest anatomical correspondence based on quantitative metrics (top row of Fig. 4), shared extensive overlap, especially in the superior parts of the TRN. In contrast, the TRNs of sub-8 and sub-10 (third row of Fig. 4) barely overlapped and were situated next to each other, with the segmentation of sub-8 being much smaller and more central than that of sub-10. Interestingly, sub-8 bore the lowest overall similarity to other subjects, with Dice coefficients ranging from 0.175 to 0.338 and AHD values ranging from 0.835 to 2.016 mm, including the highest AHD observed in the *in vivo* cohort.

**Figure 4.**
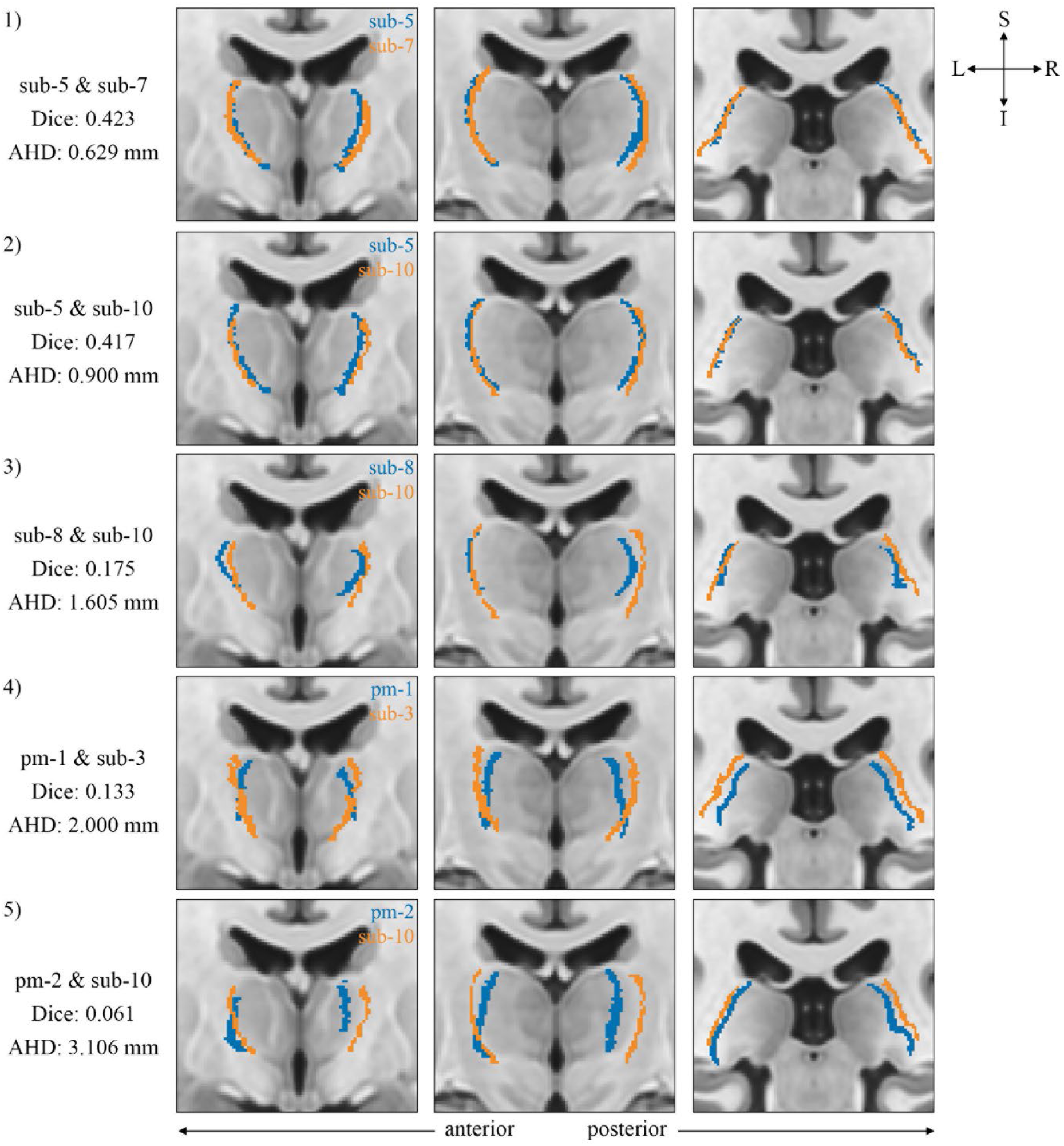
Inter-individual variability in TRN shape and overlap. Representative pairs of TRN segmentations are visualised on UNI T_1_ coronal slices in MNI space. (1) Segmentations of sub-5 (blue) and sub-7 (orange) as an example of high overlap and shape alignment, supported by the highest Dice coefficient and the lowest AHD in the sample. (2) Segmentations of sub-5 (blue) and sub-10 (orange) as an example of high overlap and moderate shape alignment. (3) Segmentations of sub-8 (blue) and sub-10 (orange) as an example of low overlap and shape alignment among *in vivo* segmentations. (4) Segmentations of pm-2 (blue) and sub-3 (orange) as an example of relatively high overlap and shape alignment between *in vivo* and postmortem-based segmentations. (5) Segmentations of pm-1 (blue) and sub-10 (orange) as an example of low overlap and shape alignment between *in vivo* and postmortem-based segmentations. Across examples, high and low correspondence are defined relative to the distribution of Dice and AHD scores within the sample. All images are coronal views, ordered from anterior (right) to posterior (left) positions. L = left; R = right; S = superior; I = inferior.

### 3.6. Comparison of the *in vivo* and postmortem TRN segmentations

The spatial overlap between *in vivo* and postmortem TRN segmentations was limited, as reflected by lower Dice scores (mean [SD] = 0.091 [0.036]) and higher AHD values (mean [SD] = 2.424 [0.352] mm). The highest Dice coefficient reached 0.156 for sub-4 and pm-2, while the lowest AHD was 1.766 mm for sub-3 and pm-2. Conversely, sub-2 and pm-1 yielded the lowest Dice coefficient (0.040) and sub-10 and pm-2 exhibited the highest AHD (3.106 mm). Qualitative inspection of the segmentation pairs indicated that these low overlap values were mainly driven by differences in spatial position rather than by differences in TRN morphology. That is, the postmortem segmentations were consistently located more medially than the *in vivo* segmentations (row 4 and 5 of Fig. 4), which substantially reduced spatial overlap. Importantly, despite the spatial offsets, the overall shape of the TRN was highly comparable across *in vivo* and postmortem data. In both modalities, the TRN appeared more elongated in superior and rostral regions and more curled in caudal regions. Accordingly, the relatively high AHD scores observed in the analysis likely reflected the differences in spatial location, rather than discrepancies in shape.

### 3.7. Probabilistic atlas of the TRN

Fig. 5 shows the probabilistic atlas of the human TRN, generated by aggregating all *in vivo* data. Voxel intensities represent the number of participants in whom this voxel was part of the TRN, with brighter colours indicating higher overlap across participants. Across both axial and coronal planes, the highest overlap occurred in the central portion of the TRN, whereas more variability was evident in the anterior and posterior parts, consistent with the inter-individual differences observed in the pairwise analyses. The resulting probabilistic TRN atlas is made publicly available to facilitate reuse in future analyses (see Data Availability section).

**Figure 5.**
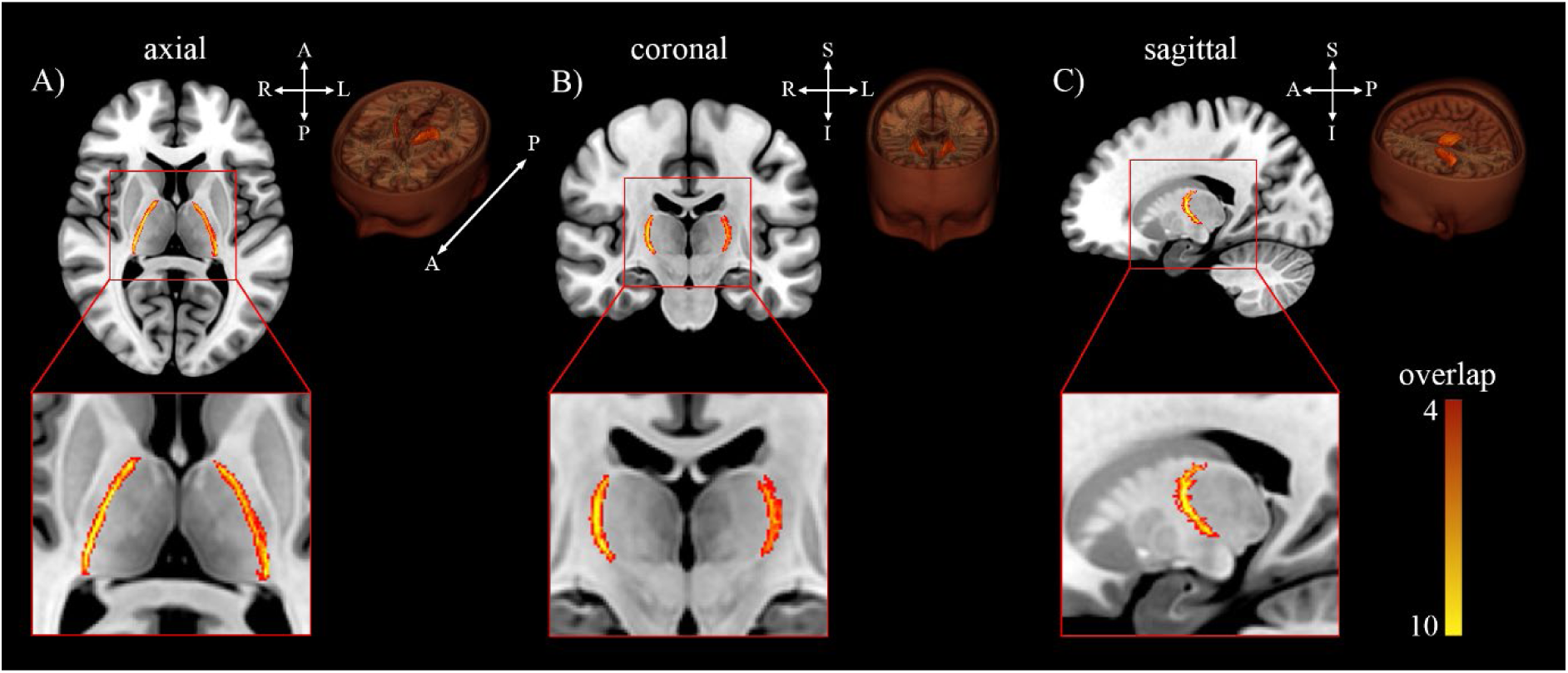
Probabilistic atlas of the human TRN. Probabilistic *in vivo* atlas of the human thalamic reticular nucleus (TRN) shown in MNI space. Voxel intensities represent the number of subjects in whom a given voxel was identified as part of the TRN, with brighter colours indicating higher inter-individual overlap. L = left; R = right; A = anterior; P = posterior; S = superior; I = inferior.

## 4. Discussion

This study assessed the feasibility of reliably identifying the human thalamic reticular nucleus (TRN) *in vivo* using ultra-high field (UHF) 7 Tesla (7T) magnetic resonance imaging (MRI) in healthy young individuals. The results provide multiple, converging lines of evidence that the human TRN can be visualised *in vivo* using high-resolution MRI. In addition, we introduce a probabilistic atlas of the human TRN, representing an important first step towards investigating its role in human brain function and disease.

Several findings support the interpretation that the delineated structure corresponds to the TRN. First, the appearance of the segmented structure was consistent with known TRN anatomy (Niemann et al., 2000; Pinault, 2004; Segobin et al., 2024) and with the postmortem datasets, where the TRN was clearly identifiable. Across datasets, the TRN appeared as a thin, elongated structure encapsulating the lateral and anterior aspects of the thalamus and bordered by the internal capsule. As expected, its appearance varied systematically with the plane and level of section, appearing more compact rostrally and superiorly and increasingly elongated and curled in caudal regions.

Second, morphometric properties extracted from the *in vivo* segmentations were biologically plausible (Niemann et al., 2000; Saranathan et al., 2021; Viviano & Schneider, 2015) and internally consistent. TRN volumes and thicknesses were stable across datasets, free of outliers, and showed no hemispheric differences. Moreover, *in vivo* estimates closely matched postmortem measurements, with small systematic differences in absolute volume likely reflecting age-related shrinkage in the postmortem sample (Tullo et al., 2019; Van Der Werf et al., 2001).

Third, although within-dataset analyses of hemispheric symmetry revealed only moderate volumetric overlap, this degree of overlap is substantial given the TRN’s thin and curved geometry. For such a structure, even small left-right asymmetries can substantially lower Dice overlap values. Importantly, the observed low AHD values indicate that hemispheric differences were driven primarily by minor spatial shifts rather than gross morphological discrepancies. Visual inspection confirmed that cases with lower overlap reflected genuine anatomical asymmetries rather than segmentation errors. Between-subject comparisons showed greater variability than within-dataset analyses, again confirmed to reflect real inter-individual differences in TRN position and extent rather than segmentation error. Importantly, spatial convergence across segmentations was observed in the probabilistic TRN atlas, with highest overlap in the central portion of the TRN and increased variability toward its anterior and posterior ends.

Finally, comparisons between *in vivo* and postmortem segmentations further supported the validity of the TRN delineations. Although spatial overlap between the two data types was limited, as reflected by low Dice coefficients, the overall TRN shape and curvature were highly consistent across modalities. The observed discrepancies were mainly attributable to systematic spatial offsets, with postmortem TRN segmentations positioned medially relative to the *in vivo* TRN. Postmortem TRN segmentations additionally appeared more asymmetrical in the axial plane and less rounded in the coronal plane, an effect that was most pronounced in pm-2. These differences are consistent with known limitations of postmortem imaging, which can alter anatomical proportions or reduce shape integrity (Schulz et al., 2011), as well as with tissue shrinkage associated with aging (Tullo et al., 2019) and postmortem tissue processing (Schulz et al., 2011).

Taken together, these results demonstrate that the human TRN is not only detectable but can be systematically characterised *in vivo* using high-resolution MRI. These findings are particularly noteworthy given that most existing anatomical knowledge of the human TRN is derived from postmortem or histological sources, which are limited in several important aspects when compared to *in vivo* imaging. While histological atlases of the thalamus offer microscopic resolution, they are often based on datasets that are distorted due to tissue shrinkage and slicing. Moreover, they typically rely on small samples, which prevents the possibility of assessing inter-individual anatomical variability. In contrast, imaging the TRN *in vivo* not only preserves anatomical integrity but also represents a first step towards studying TRN function. Ultra-high field functional MRI, in particular, offers the spatial resolution required to study TRN-related responses, and may enable future investigations of its functional role across a range of tasks and states of consciousness.

However, successfully visualising the TRN *in vivo* poses methodological challenges, as distinguishing this thin structure from surrounding nuclei requires exceptionally high spatial resolution and contrast. Ultra-high field MRI at 7 Tesla offers a significant advantage in this regard (Saranathan et al., 2021), which in this study was further strengthened by using acquisition protocols specifically tailored to this goal. The multi-echo ME GRE sequence, particularly sensitive to iron- and myelin-rich tissue (Gulban et al., 2022), proved important for defining the outer boundary of the TRN. On the other hand, data from the MP2RAGE sequence enabled clear visualisation of the thalamus, serving as a reliable reference for the inner boundary of the TRN. Repeated acquisitions of these scans allowed for image averaging during preprocessing, which enhanced the SNR and allowed for mitigation of flow-related artefacts. Together, these methodological choices were important for achieving sufficient image quality to enable reliable manual delineation of the TRN *in vivo*.

At the same time, these requirements highlight important practical limitations. The optimised protocol required long acquisition times (±1 hour), and the high spatial resolution made the data susceptible to participant motion, resulting in the exclusion of several datasets. However, note that newer sequences leveraging 3D EPI developments (Stirnberg et al., 2024) offer much faster T2*w imaging (below 7 minutes per run) with whole brain coverage at the same 0.35 mm isotropic resolution (Gulban et al., 2026), together with prospective motion corrections to mitigate within-run head motion artefacts (Serger et al., 2025). Nevertheless, while the number of available 7T scanners increases rapidly worldwide, access to 7T MRI systems remains limited compared with 3T scanners. Finally, the processing and manual segmentation are very time-intensive. As such, the approach presented here is best suited for targeted TRN studies, and not feasible as an add-on to broader neuroimaging protocols or routine clinical workflows.

Given these methodological demands and practical constraints, widespread application of individualised TRN segmentation remains challenging. To address this limitation and facilitate broader use of TRN anatomy in human neuroimaging, we provide a probabilistic *in vivo* atlas of the human TRN in the standard MNI space. This atlas can be aligned to individual datasets acquired at lower field strengths, enabling the definition of anatomically-informed regions of interest for task-based, resting-state, or connectivity analyses. When applying the atlas, careful evaluation of spatial alignment is essential. The TRN region of interest should appropriately surround the thalamus on lower resolution scans (e.g., T_1_ weighted images). Normalisation and/or alignment procedures should be refined if this anatomical correspondence is not observed.

By providing probabilistic information on the TRN’s location and shape, the atlas facilitates the integration of TRN anatomy into large-scale or lower-field neuroimaging studies without requiring manual TRN segmentation in every individual dataset. This, in turn, enables more systematic investigation of the TRN’s role in thalamocortical communication across cognitive states and experimental paradigms. Importantly, the availability of an *in vivo* TRN atlas also has clear implications for clinical research. Growing evidence links TRN dysfunction to neuropsychiatric and neurological conditions such as schizophrenia and tinnitus, and postmortem work has demonstrated marked TRN volume reduction in Alzheimer’s disease (Xuereb et al., 1991). The present atlas represents a probabilistic reference derived from healthy individuals, which could be used as normative baseline against which TRN segmentations obtained in independent or clinical populations can be compared. Such comparisons can support the identification of disease-related alterations in TRN morphology, or for example assess treatment-related changes. Extending this framework to include age-stratified atlases will be particularly important (Miletić et al., 2022), given the central role of the TRN in cognitive control and sensory filtering, processes known to change with age.

In summary, this work establishes the feasibility of delineating the human TRN *in vivo* and provides an anatomical reference for future investigations. By enabling systematic characterisation of this structure in the intact human brain, the present study bridges a critical gap between postmortem anatomy and *in vivo* neuroimaging. As methodological advances continue to improve spatial resolution and accessibility, the combination of high-resolution imaging and probabilistic atlases such as the one introduced here will be instrumental in advancing our understanding of TRN structure, variability, and its involvement in human cognition and disease.

## Author Contributions (CRediT)

**Zuzanna Kotwicka:** Data curation, Formal analysis, Methodology, Visualisation, Writing - original draft. **Omer Faruk Gulban:** Methodology, Resources, Software, Visualisation, Writing - review & editing. **Logan Dowdle:** Software, Writing - review & editing. **Ryszard Auksztulewicz:** Methodology, Supervision, Writing - review & editing. **Michelle Moerel:** Conceptualization, Funding acquisition, Investigation, Methodology, Supervision, Visualisation, Writing - review & editing.

## Data and Code Availability

All raw and derived data generated for this manuscript are publicly available on Zenodo (https://doi.org/10.5281/zenodo.20446658). This includes raw MRI data for sub-1 to sub-5 and sub-10, as well as derived files including segmentations for all datasets (sub-1 to sub-10 and pm-1 to pm-2). Raw MRI data for sub-6 to sub-9 can be accessed through the original publication of the dataset ((Gulban et al., 2022), https://doi.org/10.17605/OSF.IO/N5BJ7). Postmortem data are available from their original source (see Table 1 in (Alkemade et al., 2022)). Data analysis scripts are available on GitHub (https://github.com/mmoerel/TRN-atlas).

## Acknowledgements

We thank Alina Schepers, Sonia Bǎlan, and Sydney Kissel for their contributions to the project.

## Supplementary material

**Supplementary Figure 1.**
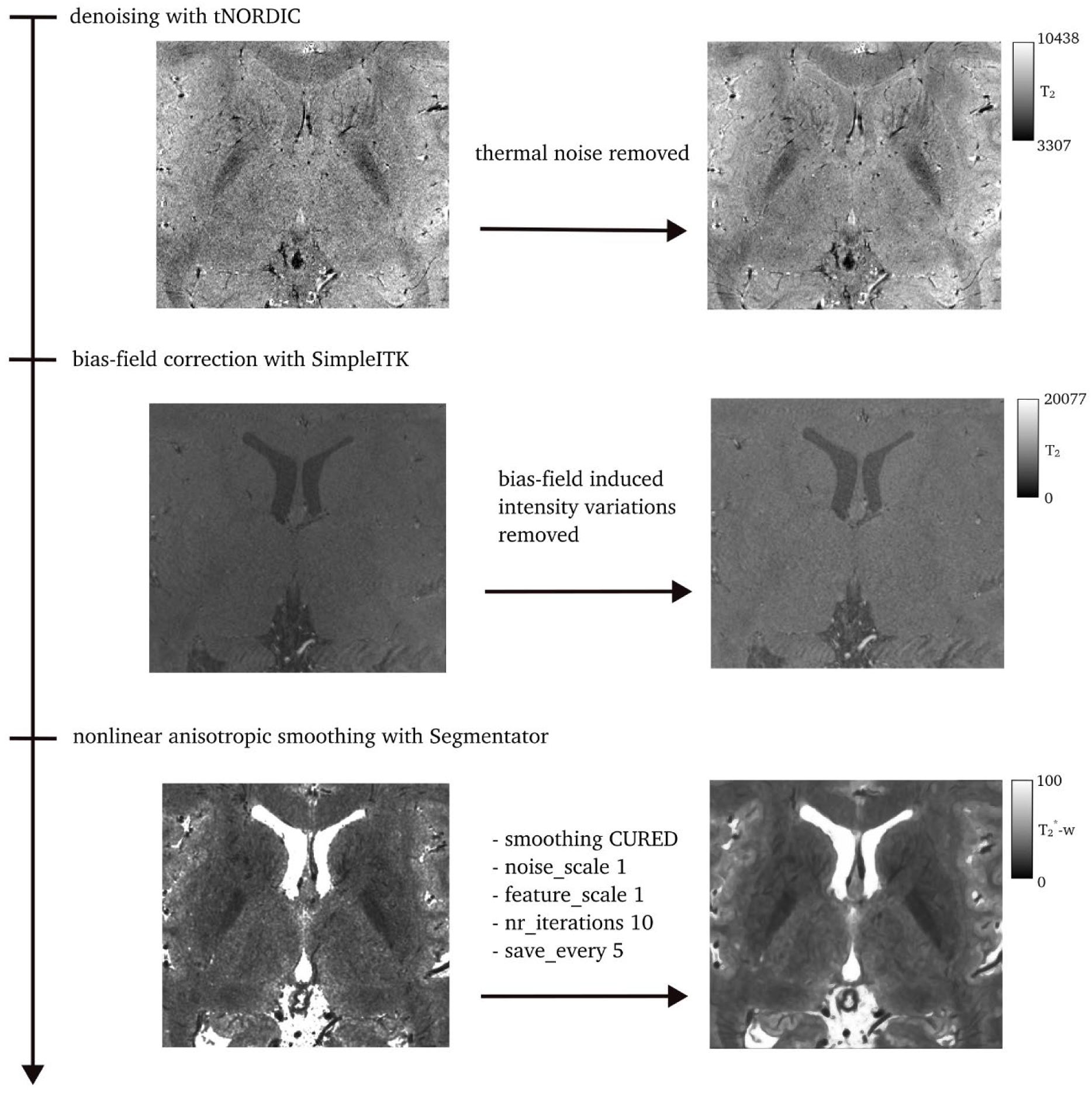
Intermediate output of the preprocessing pipeline. Output of the preprocessing steps applied to the ME GRE data prior to segmentation are shown from top to bottom. For each step, the image before processing is shown on the left and the corresponding result after processing on the right. Top row: thermal noise removal using tNORDIC. Middle row: bias-field correction performed with SimpleITK. Bottom row: nonlinear anisotropic smoothing using Segmentator.

**Supplementary Video 1.** Three-dimensional rendering of the TRN segmentation from a representative *in vivo* dataset (sub-09). The video is available at Zenodo: https://doi.org/10.5281/zenodo.20446658.

**Supplementary Figure 2.**
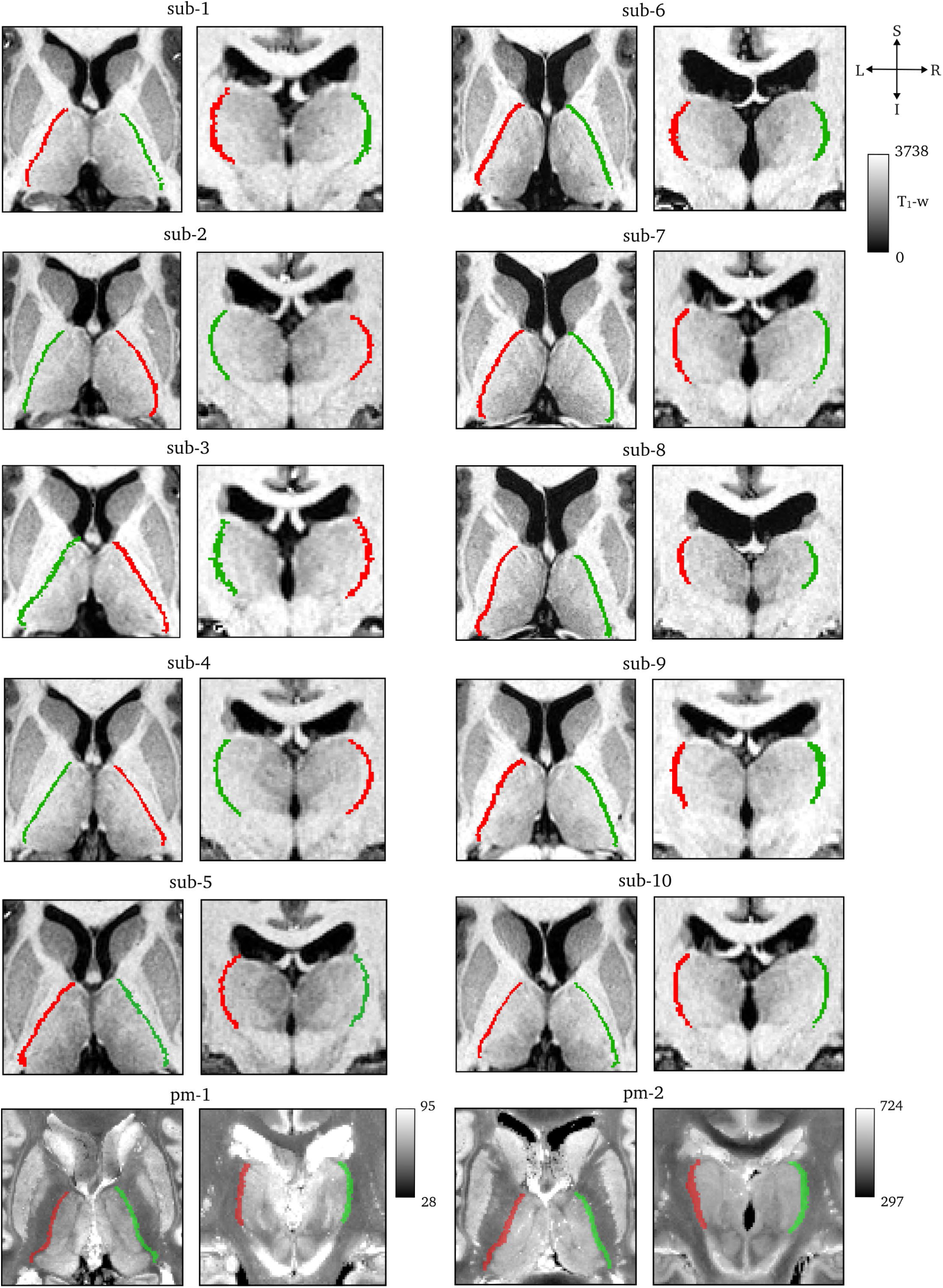
Individual TRN segmentations. Visualisation of each TRN segmentation on T_1_w MRI (sub-01 – sub-10) or blockface images (pm-1 – pm-2). For each dataset, axial views are shown on the left and coronal views on the right. S = superior, I = inferior, R = right, L = left.

